# Figure-ground segmentation based on motion in the archerfish

**DOI:** 10.1101/2022.12.25.521891

**Authors:** Svetlana Volotsky, Ronen Segev

## Abstract

Object detection and recognition is a complex computational task that is thought to rely critically on the ability to segment an object from the background. Mammals exhibit varying figure-ground segmentation capabilities, ranging from primates that can perform well on figure-ground segmentation tasks to rodents that perform poorly. To explore figure-ground segmentation capabilities in teleost fish, we studied how the archerfish, an expert visual hunter, performs figure-ground segmentation. We trained archerfish to discriminate foreground objects from the background, where the figures were defined by motion as well as by discontinuities in intensity and texture. Specifically, the figures were defined by grating, naturalistic texture, and random noise moving in counterphase with the background. The archerfish performed the task well and could distinguish between all three types of figures and grounds. Their performance was comparable to that of primates and outperformed rodents. These findings suggest the existence of a complex visual process in the archerfish visual system that enables the delineation of figures as distinct from backgrounds, and provide insights into object recognition in this animal.

## Introduction

Many animals’ visual interaction with the world relies heavily on the ability to detect and recognize objects in the environment. It is believed that surface-segmentation; i.e., the ability to delineate an object from the surrounding background, is one of the first critical stages in visual scene analysis. Thus, to better understand the computational basis of vision in different animals, more must be known about how this information processing component performs tasks.

The key visual features that identify the border of an object or surface are the discontinuities or abrupt changes at the figure border (Lamme, Victor AF and Roelfsema. 2000; Lamme, V. A. 1995; Zhou et al. 2000); i.e., changes in the properties of the surface such as luminance, orientation, or texture (Fig. 1 a). However, there are also discontinuities in these visual features in the interior of objects or surfaces. This makes the detection of actual borders using ambiguous cues a challenge to any visual system.

**Fig. 1.**
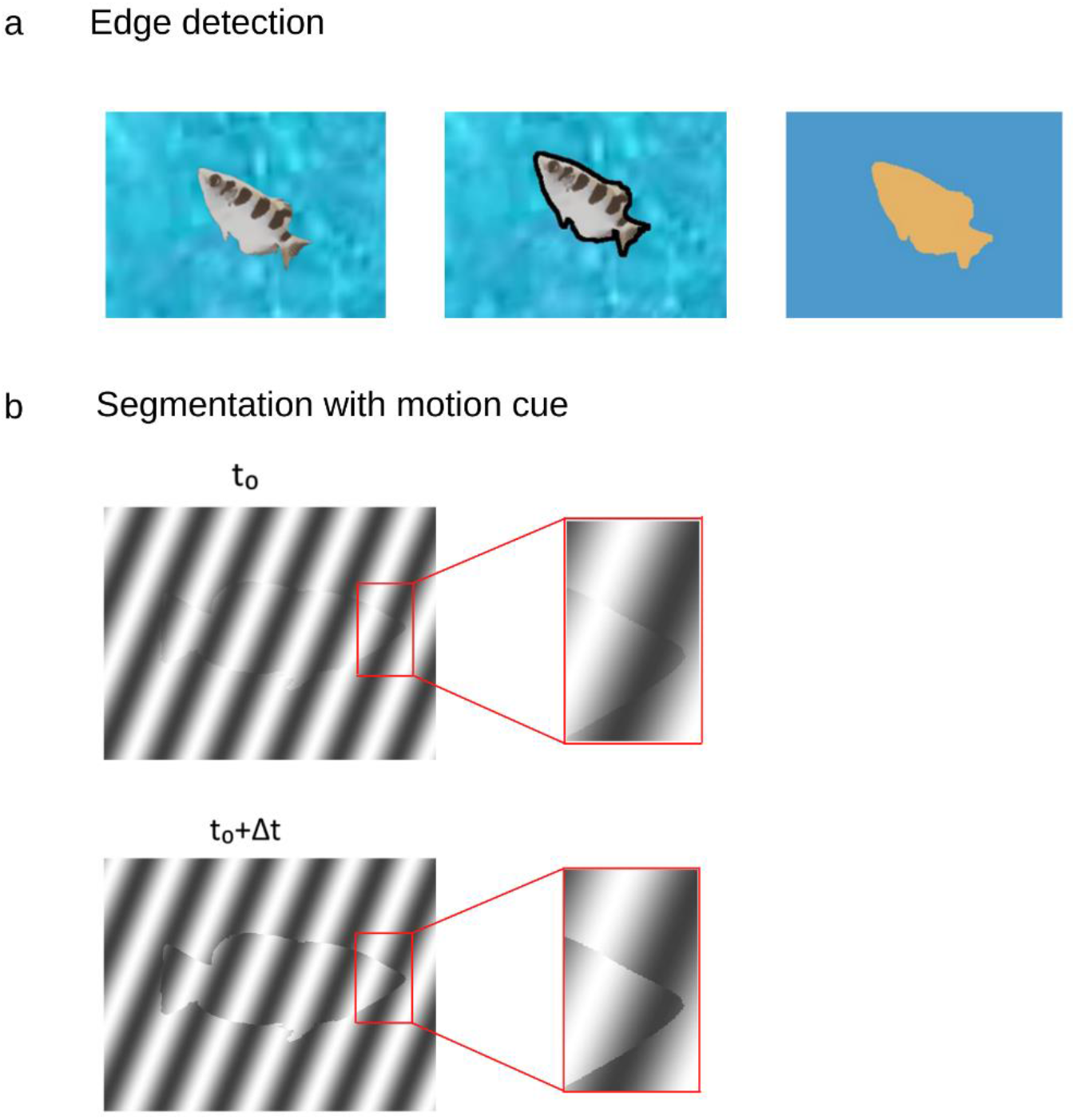
Segmentation is a crucial step in visual scene processing and object recognition. **A**. Figure-ground segmentation can be performed by edge detection based on discontinuity of intensity or color. **B**. Motion cues can be used to detect the boundary of the foreground by deletion-accretion of pixels due to the motion of the foreground over the background. By comparing the deletion and appearance of pixels resulting from object motion, the foreground can be segmented from the background.

Two cues signaling object borders are however unambiguous; namely, accretion and deletion: which are constituted by the appearance or disappearance of pixels forming the background surface due to motion of the foreground object (Fig. 1 b) (Lamme, V. A. 1995; Stoner and Albright. 1996). These are the only cues that remain unambiguous, invariant to texture and independent of learning and do not require explicit object recognition. These cues exist locally and do not require global processing of the visual scene (Gibson. 2014; Tsao and Tsao. 2022).

Behaviorally, primates’ ability to detect object borders based on accretion and deletion has been tested in several species differs considerably from that of rodents (Luongo et al. 2021; Schnabel et al. 2018). Several primate species can perform segmentation based on motion signal, both when the stimulus was based on grating, when local orientation information was present, in cases where the figure and ground were derived from naturalistic noise images, and where there was no orientation information present (Ho et al. 2021; Luongo et al. 2021; Mustafar et al. 2018).

Several rodent species were found to have limited capacity to perform surface segmentation based on accretion and deletion alone. Rodents were able to perform the figure-ground segmentation task when local orientation information could be used to segment the figure from the ground but failed to do so when the stimulus was derived from naturalistic noise images where no orientation information could be tapped (Luongo et al. 2021).

These differences between primates and rodents indicate that the visual system of different lineages perform very differently on the surface segmentation task which is so central to our understanding of visual scene processing. Clearly, to better grasp whether and how surface segmentation is performed across vertebrates’ visual systems, other species should be tested.

This article reports on teleost fish’ capacity to segment the foreground surface from the background surface. We capitalized on our ability to perform controlled experiments in the archerfish (*Toxotes Chatareus*). This fish is known for its ability to shoot insects above the water level for food (Ben-Tov et al. 2018; Newport and Schuster. 2020; Lüling. 1963; Ben-Simon et al. 2012). The archerfish can be trained to shoot at targets on a computer monitor, which enables controlled experiments. In recent years, studies have shown that the archerfish can recognize natural (Volotsky et al. 2022; Newport et al. 2016) and artificial (Newport et al. 2015; Newport et al. 2014) objects. In addition, recording of single cells in the early visual system of the archerfish (Reichenthal et al. 2018; Ben-Tov et al. 2013) showed that it has characteristics similar to those found in the early visual system of mammals, and primates in particular. This includes tuning to orientation of surfaces and contextual modulations which can be used for early processing in segmenting (Ben-Tov et al. 2013; Ben-Tov et al. 2015). Thus, the archerfish provides an excellent opportunity to study the mechanisms of object recognition in fish.

As described below archerfish were administered several segmentation tasks involving different background textures with and without a motion cue. The results showed that the archerfish can perform segmentation tasks that are purely defined by motion; i.e., the accretion and deletion of pixels. In addition, the archerfish successfully discriminated foreground objects from the background with a static target based on the discontinuity of texture. We compare these archerfish results to several mammalian species to better understand this behavior across taxa.

## Methods

### Animals

Six archerfish were used in the experiments. Adult fish (6-14 cm in length; 10-18 gm) were purchased from a local supplier. Throughout the experiments, the fish were kept separately in 100-liter aquaria filled with brackish water at 25°-29°C on a 12-12 hour light-dark cycle. Fish care and experimental procedures were approved by the Ben-Gurion University of the Negev Institutional Animal Care and Use Committee and were in accordance with government regulations of the State of Israel.

### Training

After arrival at the lab and a period of acclimatization to the laboratory conditions, the fish were trained to shoot at targets presented on a computer screen (VW2245-T, 21.5”, BenQ, Taiwan) situated 35±2 cm above the water level (Fig. 2 a). Each training session consisted of 20 trials and was conducted 2-3 times a week. First, the naïve fish were trained to shoot at a single square presented on a white background at two different locations: either in the middle of the left half of the screen or in the middle of the right half of the screen. If the fish hit the target square within 20 seconds after its appearance, it was rewarded with a food pellet. After 20 seconds the target disappeared, and the next trial started. The fish was considered ready for the experiments if it succeeded in hitting 80% of the targets in three consecutive sessions.

**Fig. 2.**
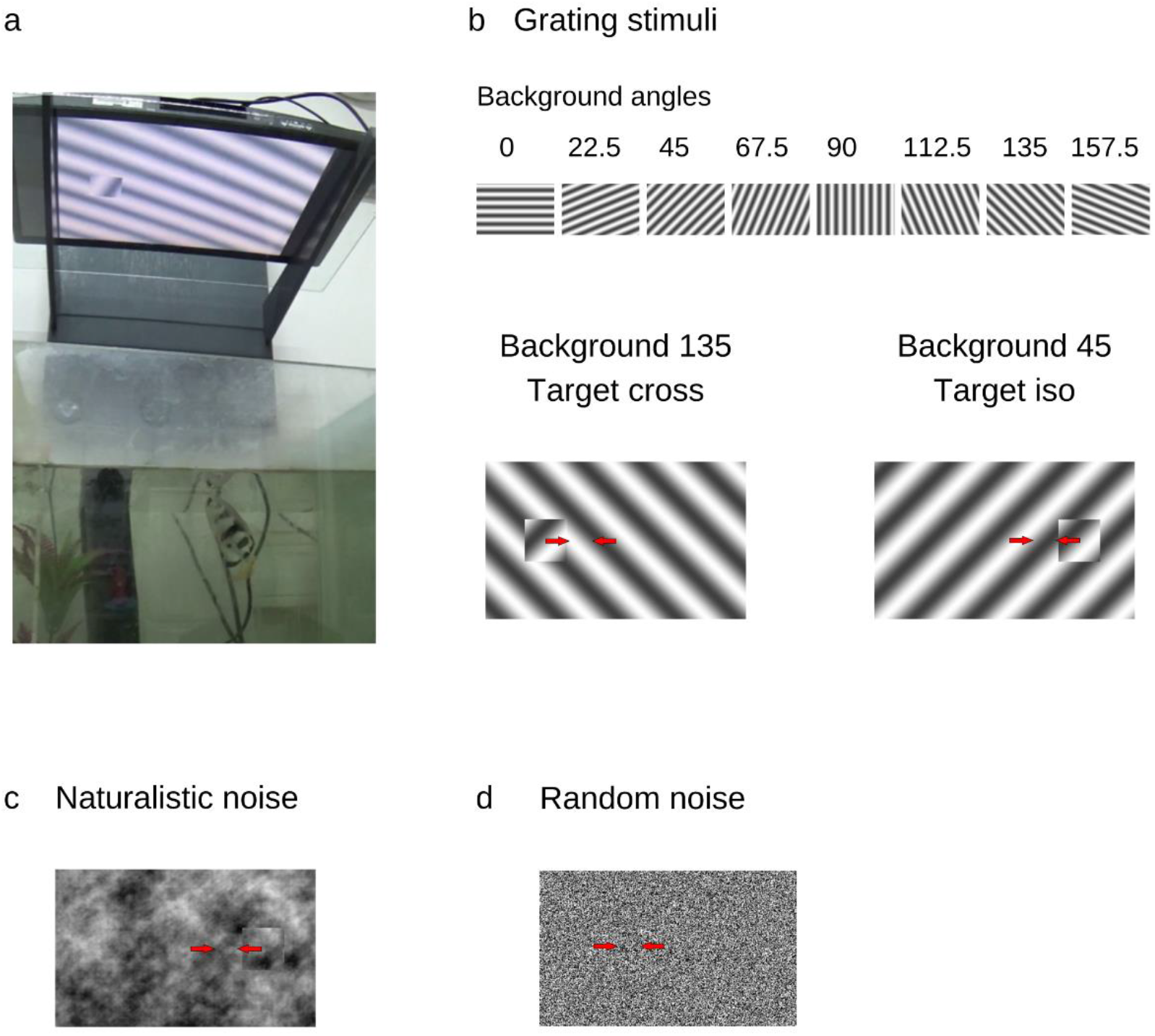
Experimental setup and different stimuli. **A**. Example of archerfish during the behavioral experiments. A screen is placed above water level and the stimulus is presented. If the fish shoots at the target, it is rewarded by placing a food pellet in the water. **B**. Grating experiment: the background orientation is selected from eight different orientations. The target is oriented either 90° from the background (Cross condition) or in the same orientation as the background (Iso condition). Arrows indicate the movement of the target and the background in the opposite directions. **C**. Example of naturalistic noise used for generating the background and the object. **D**. Example of the random noise experiment. The target boundaries vanish in the absence of the motion signal.

### Stimuli

Stimulus images were created using Matlab. For all experiments, the stimuli consisted of a surface image shown in the background and a target square image that was cropped from a surface image. The target was placed in the middle of either the left or right half of a background image. The background image measured 26 cm x 36 cm, and the target square measured 4.5 cm x 4.5 cm.

We used three types of surface textures: sinusoidal grating, naturalistic noise patterns and random noise patterns.

#### Grating

A sinusoidal grating was presented in one of 8 orientations: 0, 22.5, 45, 67.5, 90, 112.5, 135 and 157.5 degrees. The orientation of the target was either orthogonal relative to the background in the ‘cross’ condition, or parallel to the background in the ‘Iso’ condition (Fig. 2 b).

#### Naturalistic noise

To create the naturalistic noise texture, we used 200 natural images (https://www.kaggle.com/datasets/arnaud58/landscape-pictures). The images were transformed into the frequency domain. Then, the phase was randomized such that the total power stayed constant. Finally, using an inverse Fourier transform, we obtained an image in spatial domain (Fig. 2 c).

#### Random noise

The images consisted of black and white pixels. We created 200 matrices of zeros and ones that were generated with equal probability of getting zero and one in every pixel (Fig 2 d).

### Surface segmentation with the motion cue

In the first experiment we examined the ability of the fish to discriminate the target from the background with a motion cue. In each trial, the target moved 2 cm to the right and 2 cm to the left at a frequency of 1Hz. The background moved at the same speed in the direction opposite to the target. After 20 seconds of back and forth movement, the trial ended. For the grating pattern, all the possible combinations of background angle, target angle relative to the background, and target side position were used in random order. For the Naturalistic noise and Random conditions, all 200 patterns were used, such that each pattern was novel for the fish.

### Surface segmentation without the motion cue

In the second experiment we examined the ability of the fish to discriminate the target from the background without a motion cue. We used the same patterns as in the previous experiment but removed the motion such that both the target and the background were static. The position of the target was moved 1 cm from the point where the target was cropped from the texture – the farthest position of the target relative to the background. This experiment was conducted only for the grating and natural noise textures. Random noise textures were omitted because in the static condition the target would not have been visible in this case.

### Trained stimuli vs. novel stimuli

We tested whether the performance of the fish improved with training. For this purpose, the same 4 patterns of naturalistic noise targets and backgrounds were used repeatedly in 10 consecutive sessions in random order, with 5 occurrences of each pattern in every session. We examined the success rate of the fish in response to these patterns to assess improvement in their reactions to familiar patterns as compared to their responses to the novel patterns.

### Statistical analysis

All the statistical analyses were performed in Matlab. To verify the ability of the fish to discriminate the target from the background, we used the binomial cumulative distribution to estimate the target selection rate and compared it to the chance value of 50% using a binomial test. To compare the responses of the fish to the two experiments, for example, with and without a motion cue, we calculated the binomial probabilities of the fish’s success in the two experiments. If the difference between the probabilities was less than 5% on either side, the results of these two experiments were considered to be equivalent. In all statistical tests, the significance level was set at p= 0.01.

To evaluate the learning rate of the fish throughout the experiments, we estimated the slope coefficient of the linear model fitted to the fish’s success rate in separate sessions and tested whether it was significantly above zero. If the slope coefficient was significant, the change in success rate was considered to be significant.

## Results

We characterized the archerfish’s ability to perform segmentation of figure from background based on deletion-accretion and the discontinuity signal with and without a motion cue. For this purpose, we conducted two alternative non-forced choice experiments using a continuous reinforcement schedule for correct responses. The fish was presented with stimulus of a figure over a background on a computer monitor situated above the water tank. The figure was a square moving back and forth in the opposite phase over a moving background that provided a differential motion cue with accretion-deletion. The amplitude of the figure and background motion was the same (see Methods). The fish was required to find the target and shoot at it (Fig. 2 a). A shooting response directed at the figure was rewarded with a food pellet whereas shooting elsewhere was not rewarded and counted as an incorrect response. An incorrect response resulted in the termination of the trial and the initiation of the next one. To neutralize the effect of position bias in the fish’s responses, the figure was presented at a random in two locations on the screen.

We tested archerfish segmentation behavior using four classes of stimuli: 1. A cross grating stimulus, where the figure consisted of a grating and the ground consisted of an orthogonal grating (Cross condition, Fig. 2 b). 2. A grating stimulus where the figure consisted of a grating, and the ground consisted of a grating in the same orientation, but with an offset in phase (Iso condition, Fig. 2 b). The background orientation was selected at random from 8 angle options (Background angles, Fig. 2 b). 3. Naturalistic stimuli, where both the figure and the ground consisted of naturalistic noise patterns (Naturalistic condition, Fig. 2 c, see methods). 4. A random checkerboard noise where each pixel was selected randomly from a binary distribution (Random condition, Fig. 2 d).

The rationale for selecting these stimuli was motivated by the fact that grating is a classical stimulus for figure-ground segregation studies in different species (Schnabel et al. 2018; Baumann et al. 1997). In addition, the two different grating conditions (Cross and Iso) generated two similar stimuli that had different levels of difficulty associated with the task. The Naturalistic noise images capture the statistical properties of the natural images. This condition was used to differentiate between a true figure-ground signal through low level processing of orientation or a phase contrast signal (Simoncelli and Olshausen. 2001). Finally, the random checkerboard stimulus is an example of a pure accretion deletion signal with local information alone.

### Archerfish can segment objects defined by opposing motion

The results showed that the archerfish successfully performed the task based on the motion signal. The archerfish performed well from the first session of the grating stimulus presentation and maintained their nearly perfect performance thereafter (Fig. 3 a, b). This nearly perfect performance was observed in both the Cross and the Iso condition.

**Fig. 3.**
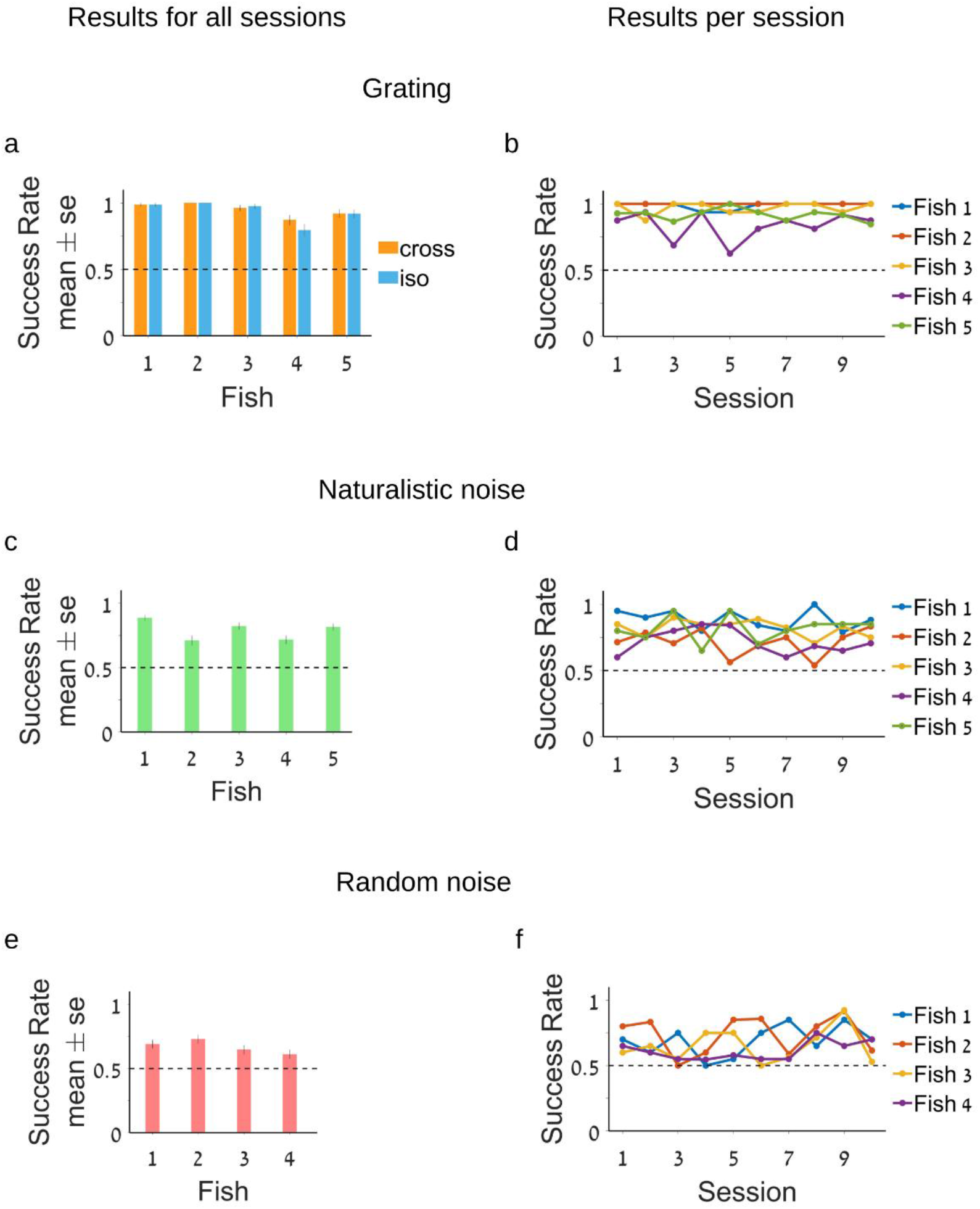
Archerfish can perform figure-ground segmentation with a motion signal. **A**. Average success rate of the fish on the segmentation task in the grating Iso and Cross conditions. **B**. Average success rate of the fish per session. The detection of the target in the Cross and Iso conditions achieved almost perfect performance. **C**. Success rate of the fish in the segmentation task in the naturalistic noise condition. **D**. Success rate of the fish per session. **E**. Random noise condition. **F**. Success rate of the fish per session. In the naturalistic noise and random noise conditions, the fish performed significantly above chance but lower than perfect performance. The fish’s success rate did not increase over sessions indicating that there was no learning in the segmentation process.

After the end of the grating experiment, the fish were presented with the Naturalistic noise stimulus, where no local orientation signal was present and thus could not be used for segmenting the figure from the ground. Again, the archerfish were able to perform the task at a success rate above chance although below their almost perfect performance in the grating experiment (Fig. 3 c, d). Finally, the fish performed surface segmentation in the extreme case of the Random condition. In this case, the figure-ground segmentation relied on the accretion-deletion signal alone. The fish were able to perform the task above chance level (p < 0.01) but with a lower success rate than the grating or natural noise (Fig. 3 e, f).

Thus overall, archerfish can segment surfaces based on the opposing signal between figure and ground. This was observed for a wide variety of stimuli but with decreasing levels of success as the task became harder.

### Archerfish segmentation in natural noise relies on the motion signal alone

Next, we examined whether the archerfish used the motion signal or some other local contrast for segmentation. For this purpose, we tested the ability of the fish to detect the same stimuli when the target was not moving, to test how critical the motion signal is in three conditions: cross grating, iso grating, and naturalistic noise. We tested the archerfish on a static condition of the task after training on the motion task.

We found that the fish were able to detect the target in both grating cases (Fig. 4 a, b). This is not surprising since in both the Cross and Iso cases, the single static frame still has an edge contrast as a result of orientation or phase differences. Thus, the image contained clear figure edges that could be detected.

**Fig. 4.**
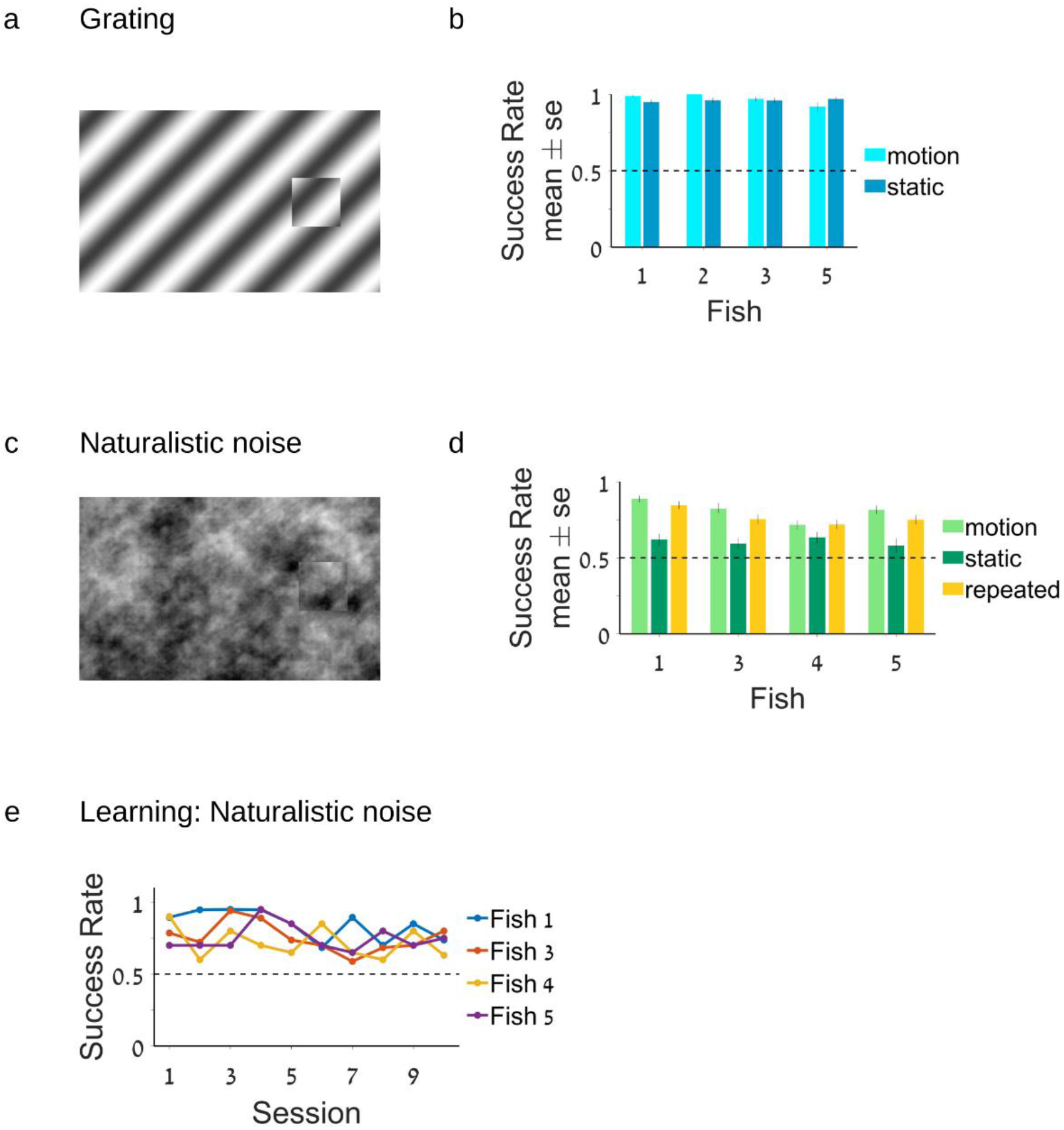
Comparison between motion cue and static cues for different stimulus types and repeated stimuli. **A**. Example of the grating stimulus. **B**. In the grating stimulus condition, the fish detects the target and performs the segmentation process in both conditions, with and without the motion cue. **C**. Example of the naturalistic noise stimulus. **D**. In the naturalistic noise case, the fish can perform segmentation using the motion cue but fail to do so in the static condition (light green vs. dark green). There was no significant difference in fish performance in the case of novel vs. repeated stimuli (light green vs. yellow) **E**. Success rate of the fish in the repeated naturalistic noise condition per session. The fish did not show any clear learning in terms of improvement over sessions.

In contrast, the fish could not detect the object in the case of static naturalistic noise. All fish, with one exception, performed near chance level and did not improve over sessions (Fig. 4 c, d, motion and static). This was expected since the discontinuity cue from the boundary is much weaker in the case of naturalistic noise than in the grating condition (see for example Fig. 4 a, c). Hence, motion is a critical component of the fish’s ability to perform figure-ground segmentation.

Overall, these experiments confirm that the archerfish used the motion signal to perform figure-ground segmentation in the natural noise case. The results for the grating experiment indicate that the fish used different strategies in the naturalistic and grating conditions. It is important to note that the random condition was not tested since in this case, there is no signal of the target position. We therefore omitted this condition from the experiment.

### Response to repeated stimuli shows that archerfish do not learn the task

Initially, we tested the ability of the archerfish to perform object segmentation on a naturalistic noise background using 200 different non-repeated stimuli. We looked at the average response of the fish and also at the change in success rate over the course of the 10 sessions of the experiment. We observed variability in the fish’s responses across different sessions that reached 30% in some fish, but no improvement over time. The average success rate in the first sessions of the experiments was similar to the success rate in the last sessions.

We then assessed whether the fish would improve their performance if we trained them on the same repeated backgrounds. We used 4 different backgrounds and presented them to the fish in 10 sessions, with 5 repetitions of each background per session. We observed no improvement in any of the 4 fish that completed the experiment (Fig. 4 e). To calculate the learning rate and test its significance, we fitted a linear model to the success rate of each fish in every session. None of slope coefficients of the fitted linear models was significant. This is an indication that learning, if it exists, must be very weak. When we compared the average success rate in fish’s responses to novel vs. repeated backgrounds (Fig. 4 d, motion and repeated), results were comparable.

### Comparing segmentation behavior in archerfish, rodents, and primates

In a recent study (Luongo et al. 2021), the segmentation behavior of four different mammalian species was studied in two primate species: – the macaque and mouse lemur, mice of the *Rodentia* order and treeshrews of the *Scandentia* order. This provided an opportunity to compare the archerfish segmentation behavior with these species since the experimental procedures were identical. Figure 5 presents a comparison of the success rate of these different species. In the grating experiments, the archerfish, like the other species, was able to perform the task in both the Iso and Cross conditions. The archerfish and the primates exhibited slightly better performance (Fig. 5 a, b). In the naturalistic noise condition, a different result emerged. The archerfish and primates could identify the target defined by an accretion-deletion signal whereas rodents performed at chance level (Fig. 5 c). Hence the archerfish figure-ground segmentation appears to be comparable to primates. Specifically, the archerfish performance can be considered to be situated between the macaque and the mouse lemur.

**Fig. 5.**
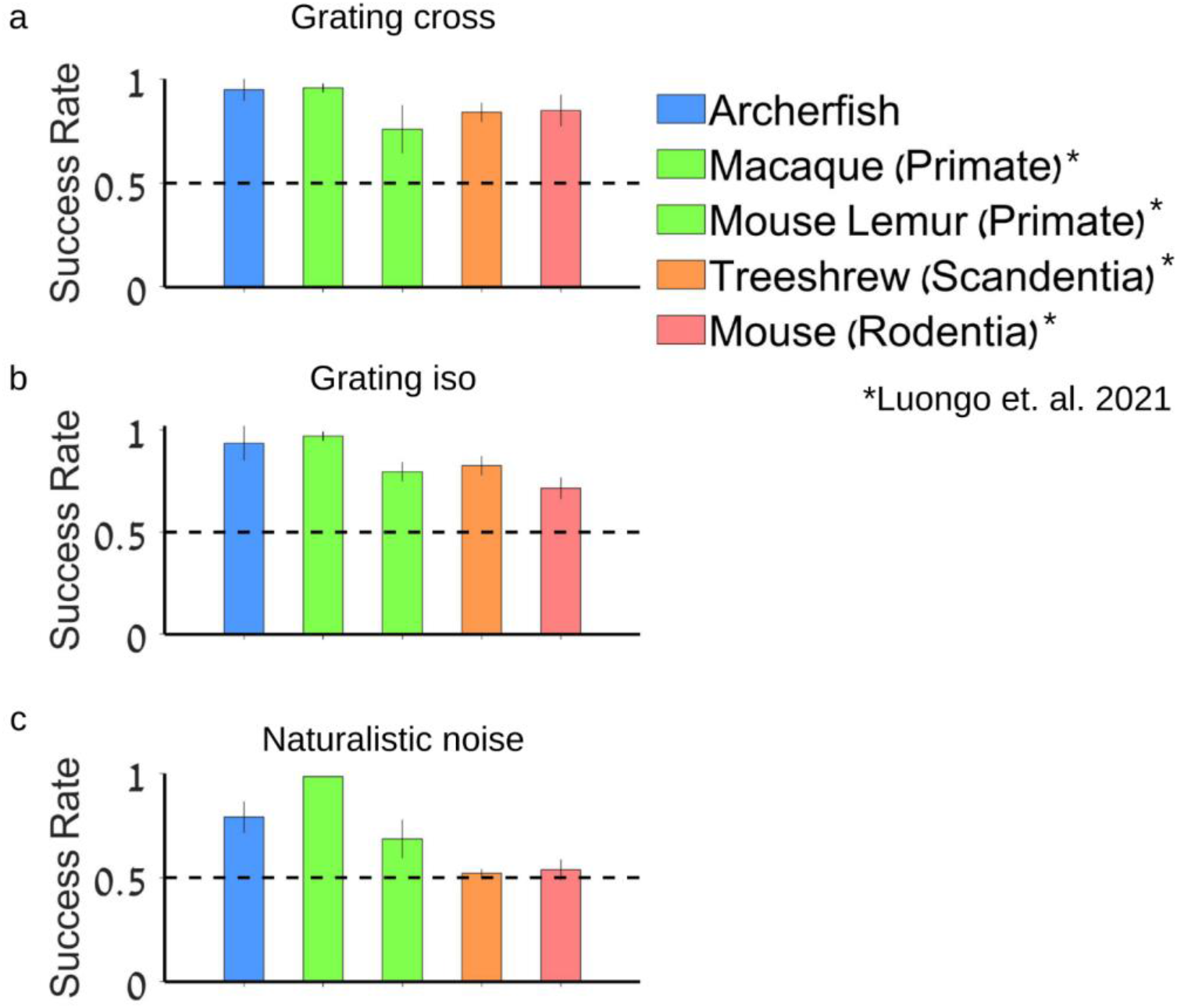
Comparison of archerfish to other species. **A**. Grating cross condition average success rate. **B**. Grating iso condition. **C**. Naturalistic noise condition. Comparison of the ability of the archerfish on different tasks indicates that the archerfish is comparable to primates since it was successful on both the grating and naturalistic noise conditions using the motion cue.

## Discussion

We studied the ability of archerfish to perform figure-ground segmentation based on different cues: accretion-deletion, discontinuity and a combination of both. In the case of accretion-deletion, the segmentation process relies on the accretion-deletion of pixels before and after the moving figure. We used grating, naturalistic noise, and random noise as targets moving counterphase with the background.

We found that the archerfish succeeded in delineating the target from the background in all types of stimuli when a motion cue was present. The fish were also able to generalize from the moving targets to the static targets, since they performed the grating task at a high success rate from the very first session. The archerfish needed the motion signal in the case of the naturalistic noise because the fish failed to detect the target in the static case.

This study draws on Loungo et al. 2021 who examined the ability of several mammalian species across three lineages to perform a figure-ground segmentation task. Their work underscored the importance of a comparative study of the mechanism of figure-ground segmentation. They found diversity in performance across mammalian lineages, which is indicative of the inherent differences and difficulties in the processing of objects across species. Specifically, they found that mice (order *Rodentia*) and treeshrew (order *Scandentia*) were less able to exploit a pure motion signal as compared to primates where the mice could not perform the naturalistic noise condition based on the motion signal. We implemented use the same stimuli to enable direct comparison across taxa.

We found that archerfish are comparable to the primates in their capacity to use a motion signal combined with discontinuity to perform figure-ground segmentation (Fig. 5). This was true in the case of their success rate under different conditions with the moving target and their inability to perform the task when the target was static.

This finding is important since it indicates that the capacity to use a motion signal to segment figure from ground has evolved independently across taxa. As fish and mammals diverged about 450 MY ago, the ability to perform segmentation based on a motion signal has probably evolved several times whenever this ability was needed.

Previous literature on figure ground segmentation in fish has only examined conditions where the visual cues consisted of intensity or orientation contrast in redtail splitfins (Sovrano and Bisazza, 2009) and in archerfish (Mokeichev et al. 2010). These works reported that fish can perceive subjective, or illusionary, contours when these contours lack a physical counterpart in terms of luminance contrast gradients. These contours can be based on either continuity with nearby real contours or changes in texture that define the contour. These studies differ from the work reported here since the segmentation process did not use motion as a cue. The present study is thus the first to focus on motion as the single unambiguous cue for figure-ground segmentation in fish.

In the archerfish, studies on object recognition generally deal with cases where the object could be easily segmented from the ground based on discontinuity of intensity. This was observed in both the case of object recognition of naturalistic objects (Volotsky et al. 2022) and in the case of human face recognition (Newport et al. 2018). In addition, in previous works, the targets did not move, although it well-known that motion is salient visual cue for the archerfish visual system (Ben-Tov et al. 2015; Reichenthal et al. 2019).

Another lineage that has received considerable attention in the literature on figure-ground segmentation is birds and especially pigeons (Lazareva et al. 2006a; Lazareva et al. 2006b). Studies on birds suggest they may attend to the figural region rather than use local properties when performing figure-ground segmentation. It was suggested that specific regions in the brain might be responsible for figure-ground segmentation (Scully et al. 2014). A recent work on the barn owl tested its ability to detect a figure using the animal’s self-motion and found that information about self-motion can facilitate orientation-based figure-ground segmentation (Dutta et al. 2020). The present study is complementary to these works since we explored segmentation in another taxon.

The study of figure ground segmentation across species raises the question what is the neural basis of this visual processing capability. As the knowledge about the function of single cells in the early visual system is still lacking in some species and exists in others (Luongo et al. 2021; Reichenthal et al. 2018; Self et al. 2013; Baumann et al. 1997) there is a need for future work to address this question in full.

## Conclusion

We examined the ability of archerfish to segment figure from ground based on motion cue. Indeed, the archerfish were able to use the motion signal, however, depending on the stimuli, the fish could use additional orientation signal for segmentation. Future studies should explore whether and how this visual segmentation process is carried out by the neural circuitry responsible for object recognition in the archerfish.

## Acknowledgments

We would like to thank Ayal Moshe Green, Ilan Rosenstein and Hadas Wardi for technical assistance.

## References

Baumann R, van der Zwan R, Peterhans E (1997) Figure-ground segregation at contours: a neural mechanism in the visual cortex of the alert monkey. Eur J Neurosci 9:1290–303

Ben-Simon A, Ben-Shahar O, Vasserman G, Ben-Tov M, Segev R (2012) Visual acuity in the archerfish: behavior, anatomy, and neurophysiology. Journal of Vision 12:18-

Ben-Tov M, Kopilevich I, Donchin O, Ben-Shahar O, Giladi C, Segev R (2013) Visual receptive field properties of cells in the optic tectum of the archer fish. J Neurophysiol 110:748–59. doi: 10.1152/jn.00094.2013 [doi]

Ben-Tov M, Ben-Shahar O, Segev R (2018) What a predator can teach us about visual processing: a lesson from the archerfish. Curr Opin Neurobiol 52:80–7

Ben-Tov M, Donchin O, Ben-Shahar O, Segev R (2015) Pop-out in visual search of moving targets in the archer fish. Nature communications 6

Dutta A, Lev-Ari T, Barzilay O, Mairon R, Wolf A, Ben-Shahar O, Gutfreund Y (2020) Selfmotion trajectories can facilitate orientation-based figure-ground segregation. J Neurophysiol 123:912–26

Gibson JJ (2014) The ecological approach to visual perception: classic edition. Psychology press

Ho CLA, Zimmermann R, Weidinger JDF, Prsa M, Schottdorf M, Merlin S, Okamoto T, Ikezoe K, Pifferi F, Aujard F (2021) Orientation preference maps in Microcebus murinus reveal size-invariant design principles in primate visual cortex. Current Biology 31:733,741.e7

Lamme VA, Roelfsema PR (2000) The distinct modes of vision offered by feedforward and recurrent processing. Trends Neurosci 23:571–9

Lamme VA (1995) The neurophysiology of figure-ground segregation in primary visual cortex. J Neurosci 15:1605–15

Lazareva OF, Castro L, Vecera SP, Wasserman EA (2006) Figure-ground assignment in pigeons: Evidence for a figural benefit. Percept Psychophys 68:711–24

Lazareva OF, Vecera SP, Wasserman EA (2006) Object discrimination in pigeons: Effects of local and global cues. Vision Res 46:1361–74

Lüling K (1963) The archer fish. Sci Am 209:100–9

Luongo FJ, Liu L, Ho CLA, Hesse JK, Wekselblatt JB, Lanfranchi F, Huber D, Tsao DY (2021) Mice and primates use distinct strategies for visual segmentation. BioRXiv

Mokeichev A, Segev R, Ben-Shahar O (2010) Orientation saliency without visual cortex and target selection in archer fish. Proc Natl Acad Sci U S A 107:16726–31. doi: 10.1073/pnas.1005446107 [doi]

Mustafar F, Harvey MA, Khani A, Arato J, Rainer G (2018) Divergent Solutions to Visual Problem Solving across Mammalian Species. eNeuro 5:10.1523/ENEURO.0167,18.2018. eCollection 2018 Jul-Aug. doi: ENEURO.0167-18.2018 [pii]

Newport C, Schuster S (2020) Archerfish vision: Visual challenges faced by a predator with a unique hunting technique. 106:53–60

Newport C, Wallis G, Reshitnyk Y, Siebeck UE (2016) Discrimination of human faces by archerfish (Toxotes chatareus). Scientific reports 6:27523

Newport C, Wallis G, Siebeck UE (2018) Object recognition in fish: accurate discrimination across novel views of an unfamiliar object category (human faces). Anim Behav 145:39–49

Newport C, Wallis G, Siebeck UE (2015) Same/different abstract concept learning by archerfish (Toxotes chatareus). PloS one 10:e0143401

Newport C, Wallis G, Siebeck UE (2014) Concept learning and the use of three common psychophysical paradigms in the archerfish (Toxotes chatareus). Frontiers in neural circuits 8:39

Reichenthal A, Ben-Tov M, Ben-Shahar O, Segev R (2019) What pops out for you pops out for fish: Four common visual features. Journal of vision 19:1-

Reichenthal A, Ben-Tov M, Segev R (2018) Coding Schemes in the Archerfish Optic Tectum. Frontiers in neural circuits 12:18

Schnabel UH, Bossens C, Lorteije JA, Self MW, Op de Beeck H, Roelfsema PR (2018) Figure-ground perception in the awake mouse and neuronal activity elicited by figure-ground stimuli in primary visual cortex. Scientific reports 8:1–14

Scully EN, Acerbo MJ, Lazareva OF (2014) Bilateral lesions of nucleus subpretectalis/interstitio-pretecto-subpretectalis (SP/IPS) selectively impair figure–ground discrimination in pigeons. Vis Neurosci 31:105–10

Self MW, van Kerkoerle T, Super H, Roelfsema PR (2013) Distinct roles of the cortical layers of area V1 in figure-ground segregation. Current biology 23:2121–9

Simoncelli EP, Olshausen BA (2001) Natural image statistics and neural representation. Annu Rev Neurosci 24:1193–216

Sovrano VA, Bisazza A (2009) Perception of subjective contours in fish. Perception 38:579–90

Stoner GR, Albright TD (1996) The interpretation of visual motion: evidence for surface segmentation mechanisms. Vision Res 36:1291–310

Tsao T, Tsao DY (2022) A topological solution to object segmentation and tracking. Proceedings of the National Academy of Sciences 119:e2204248119

Volotsky S, Ben-Shahar O, Donchin O, Segev R (2022) Recognition of natural objects in the archerfish. J Exp Biol 225:jeb243237

Zhou H, Friedman HS, Von Der Heydt R (2000) Coding of border ownership in monkey visual cortex. Journal of Neuroscience 20:6594–611

